# A novel vaccination strategy induces vaccine-specific mucosal responses at port of viral entry and exit: using systemic SARS-CoV-2 vaccination as a test case

**DOI:** 10.1101/2025.10.09.681417

**Authors:** Rupsha Fraser, Jürgen Schwarze, David Dockrell

**Author notes:** Corresponding author: Rupsha Fraser.

## Abstract

Respiratory infectious diseases are associated with substantial morbidity and mortality rates worldwide, especially at the extremes of age and in immunocompromised individuals. Over the past two decades, respiratory viruses have driven nine epidemics and pandemics (including the COVID-19 pandemic); and their ongoing emergence, persistence and evolution continue to threat global health security. The upper respiratory tract (URT) represents the primary access point for respiratory viruses, where initial host infection occurs. Vaccine-mediated URT mucosal memory responses can control infection, prevent transmission and limit viral evolution. However, vaccines against respiratory viruses predominantly have systemic administration routes that elicit strong responses in the circulation to prevent severe respiratory disease, but do not effectively block infection and onward community transmission. To overcome the limitations of systemic vaccination alone, we present a novel intervention combining systemic vaccination with targeted non-antigenic inflammatory stimulation of the URT, to induce vaccine-specific immune responses in the URT mucosa. Using SARS-CoV-2 vaccination as a test case, we demonstrate for the first time, that intranasal coadministration of exogenous IFN-α as a targeted inflammatory URT signal, alongside systemic vaccination, induces vaccine-specific T-cell responses in the URT.

## Introduction

Community-acquired respiratory infections are collectively recognised as a leading healthcare and economic burden worldwide. Vaccines remain the most promising countermeasure for mitigating future pandemic threats posed by respiratory pathogens. However, the majority of vaccines for respiratory viruses are parenterally administered, which generates systemic vaccine-specific adaptive immunity but with limited delivery to the URT mucosa, and therefore poor induction of vaccine-specific adaptive responses at the infection site ^1,2^. Indeed, nasal SARS-CoV-2-specific T-cell responses have been detected in vaccinees with SARS-CoV-2 breakthrough infections, but not in uninfected vaccinees ^3^, indicating that systemic SARS-CoV-2 vaccines alone do not prevent the acquisition of SARS-CoV-2 infection.

While intranasal vaccination platforms can elicit robust vaccine-specific URT responses in experimental animals, the vast majority of intranasal vaccines in humans have to date failed to meet efficacy criteria for success. One of the biggest challenges in developing effective intranasal vaccines in humans, is nasal mucociliary clearing, affecting drug absorption and bioavailability, with the consequential requirement for high doses of intranasal antigen delivery that exceed safety limits for human use, as discussed in Fraser et al. 2023 ^4^. To date, the only licensed intranasal vaccine is the trivalent paediatric intranasal live attenuated influenza vaccine (LAIV; Fluenz^®^in Europe, FluMist^®^in North America). In contrast, other licensed influenza vaccines are parenteral and induce systemic responses, but not URT mucosal immunity ^5^. Alternative proposed methods for induction of vaccine-specific memory responses in the URT mucosa involve systemic priming and subsequent mucosal boosting via intranasal antigen administration ^6,7^. However, the intranasal route of antigen delivery in these strategies approaches aligns with the intranasal antigen delivery in intranasal vaccines, and poses similar absorption and bioavailability challenges to intranasal vaccination.

We therefore present an innovative intervention whereby antigen-specific URT responses are induced by introducing a non-antigenic inflammatory signal that can bypass the challenges associated with intranasal vaccination/antigen administration, using SARS-CoV-2 vaccination as a test case. Our approach excludes any form of intranasal antigen administration, instead utilising coadministration of intranasal interferon (IFN)-α with systemic SARS-CoV-2 vaccination, to promote the recruitment of vaccine-induced antigen-specific lymphocytes to the URT. In this proof-of-concept study, we provide the first direct evidence that targeted induction of a non-antigenic inflammatory signal in the URT via intranasal IFN-α, with concurrent systemic vaccination, can effectively induce the localisation of vaccine-specific lymphocytes to this critical site of viral entry. This enhanced prophylactic strategy may offer a convenient solution for generating protective immunity against respiratory viral infection at the main port of viral entry and exit, thereby limiting infection and onward community transmission.

## Methods

### Study design

The goal of this study was to demonstrate the utility of intranasal IFN-α coadministration with systemic SARS-CoV-2 vaccination, in generating vaccine-specific immunity at the URT (i.e. the main port of viral entry and exit). Following treatments, T-cell responses in the URT were assessed to determine vaccine-reactivity. We emphasise that since existing licensed products can be utilised for the proposed intervention, without any necessary modification to vaccine formulation, an accelerated route to market approval and patient benefit is supported.

### Mice

Murine studies were conducted under Home Office project license number PP8738752, and in accordance with the UK Animals Act. Animals were housed in specific pathogen-free conditions in ventilated cages, at a constant temperature and humidity, and a fixed light-dark lighting cycle. Treatments were administered at the same time of day to avoid circadian rhythm effects, and animals were cohoused prior to commencement of and during experiments to normalise microbiome effects. Animals were also monitored for potential side effects of intranasal IFN-α administration, such as localised haemorrhage, anaphylaxis or >15% weight loss.

### Mouse experiments

Female C57BL/6 mice (9–12 weeks) were randomised to groups and anaesthetised with isoflurane. Animals received intraperitoneal vaccination with 6 × 10^9^ viral particles of Vaxzevria AZD1222 (AstraZeneca SARS-CoV-2 vaccine; 61µl) on days 0 and 21, with or without intranasal coadministration of 4 × 10^5^ U recombinant mouse IFN-α (4 × 10^5^ U IFN-α/10µl PBS; 5µl/nostril, 10µl total per animal) (BioLegend London, UK), and were euthanised on day 42 together with naïve controls. In an additional control experiment, mice were given 4 × 10^5^ U intranasal IFN-α alone (without concomitant systemic vaccination) on days 0 or 19 and euthanised on day 21 (21 days or 48 hours post treatment). Following euthanisation, URT tissues (nasopharynx-associated lymphoid tissue, nasal turbinates, cervical lymph nodes) were harvested, and single cell suspensions prepared for *ex vivo* stimulation with SARS-CoV-2 Spike glycoprotein peptide pools (containing the surface antigen from AZD1222). Briefly, tissues were digested with 3mg/ml Collagenase D (Roche, Hertfordshire, UK) and 30μg/ml DNase I (Thermo Fisher Scientific, Dorset UK) and processed in a GentleMACS tissue dissociator (Miltenyi Biotec, Surrey, UK). Red blood cells in the homogenates were lysed with ACK lysis buffer (Gibco, Paisley UK), and suspensions washed, centrifuged 500 x *g* for 10 minutes at 4°C, and filtered through a 70-μm cell strainer.

### *Ex vivo* cell stimulation

Single-cell suspensions were resuspended in phenol red-free RPMI 1640 supplemented with 10% FBS, 2 mM L-glutamine, 1% MEM non-essential amino acids, 1% sodium pyruvate, 0.1% 2-mercaptoethanol, 50 µg/ml penicillin/streptomycin, and 2.5 µg/ml amphotericin (Gibco, Paisley, UK). Cells were stimulated with SARS-CoV-2 Spike peptide pools (1 µg/ml per test; ImmunoServ, Cardiff, UK) for 18 h at 37°C in 5% CO_2_, with 5µg/ml brefeldin A and 2µM monensin (BioLegend, London, UK) added after 4 hours to block extracellular IFN-γ transport. Pooled cells from each group were stimulated with a non-specific stimulation cocktail (eBioscience, Altrincham, UK) under the same conditions, as a positive control.

### Flow cytometry

Increased intracellular IFN-γ expression by CD3^+^CD4^+^ or CD3^+^CD8^+^ T-cells was assessed by flow cytometry to determine specific reactivity to SARS-CoV-2 Spike peptide pools (i.e. vaccine-reactivity). Following 18-hour stimulation, cells were washed twice with PBS and centrifuged at 350 x *g* for 5 minutes at 4°C, stained with a surface marker antibody cocktail containing CD45-AF700, CD3-FITC, CD4-BV605, CD8a-PerCP/Cy5.5 antibodies (BioLegend, London, UK) for 45 minutes in the dark at RT. An additional aliquot of cells taken from a pool of all samples was kept aside as a negative control without subsequent IFN-γ staining. All cells (minus the pooled sample serving as a negative control) were washed with PBS and centrifuged at 350 x *g* for 5 minutes at 4°C, fixed and permeabilised using the Cyto-Fast™ Fix/Perm kit according to manufacturer’s instructions (BioLegend, London, UK), followed by intracellular staining using an IFN-γ-APC antibody (BioLegend, London, UK) for 45 minutes in the dark at RT. Cells were washed twice with PBS and centrifuged at 350 x *g* for 5 minutes at 4°C, and resuspended in FACS buffer for flow cytometric analyses. Sample acquisition was performed on a BD LSR Fortessa 5L flow cytometer (BD Life Sciences, Berkshire, UK) and data were analysed using FlowJo software.

### Experimental group size power calculation

Using a two-sided two-sample t-test with a significance level of 0.017, the effect size (δ) was calculated as δ = (µ_1_ - µ_2_) /σ based on the frequency of CD3+CD4+IFN-γ+ T-cells: δ = 8.086, (µ_1_ = 47.380; μ_2_ = 1.800; σ = common SD = 5.989). For the minimum detectable difference, with 80% power, an experimental group size of at least n=3 is required.

### Statistical analysis

Frequences of CD3^+^CD8^+^IFN-γ^+^and or CD3^+^CD8^+^IFN-γ^+^ T-cells were compared between experimental groups using two-way ANOVA and post hoc Tukey’s multiple comparison tests (GraphPad Prism Software, San Diego, CA, USA). *P* < 0.05 was considered statistically significant.

## Results

### Intranasal IFN-α and systemic AZD1222 coadministration induced vaccine-specific T-cell responses in the URT

To evaluate the induction of antigen-specific T-cell responses in the URT, intracellular cytokine staining and flow cytometric analysis were performed following *ex vivo* stimulation with SARS-CoV-2 Spike peptide pools. Vaccine-specific T-cells were defined as CD45^+^CD3^+^CD4^+^IFN-γ^+^ or CD45^+^CD3^+^CD8^+^IFN-γ^+^ cells, identified using sequential gating on cell size (SSC-A vs. FSC-A), singlets (FSC-H vs. FSC-A), and viability (LIVE/DEAD-negative). FlowJo-based quantification revealed a marked induction of IFN-γ-producing CD4^+^ and CD8^+^ T-cells in the URT following coadministration of intranasal IFN-α with systemic AZD1222 vaccination in C57BL/6 mice. The percentage of IFN-γ^+^CD4^+^ or IFN-γ^+^CD8^+^ T-cells in peptide-stimulated samples was calculated after subtracting background signal from the negative control (fluorescence minus one; FMO), and values were then expressed as relative to the response observed in the positive control (cells stimulated with a non-specific cell stimulation cocktail, eBioscience). IFN-γ^+^CD4^+^ T-cell frequency was significantly higher in the intranasal IFNα + systemic AZD1222 group (37.34%) compared to the systemic AZD1222 alone group (0.16%) (P < 0.001) and also compared to the untreated group (0.01%) (P < 0.001) (Figure 1). Likewise, IFN-γ^+^CD8^+^ T-cell frequency was significantly higher in the intranasal IFNα + systemic AZD1222 group (47.38%) compared to systemic AZD1222 alone (1.8%) (P < 0.001) and also compared to the untreated group (0.096%) (P < 0.001) (Figure 2). These data demonstrate that local IFN-α administration synergises with systemic AZD1222 to elicit URT mucosal T-cell immunity. Importantly, animals exhibited stable body weights and no signs of clinical distress throughout the experiment, indicating that this regimen was safe and well-tolerated.

**Figure 1.**
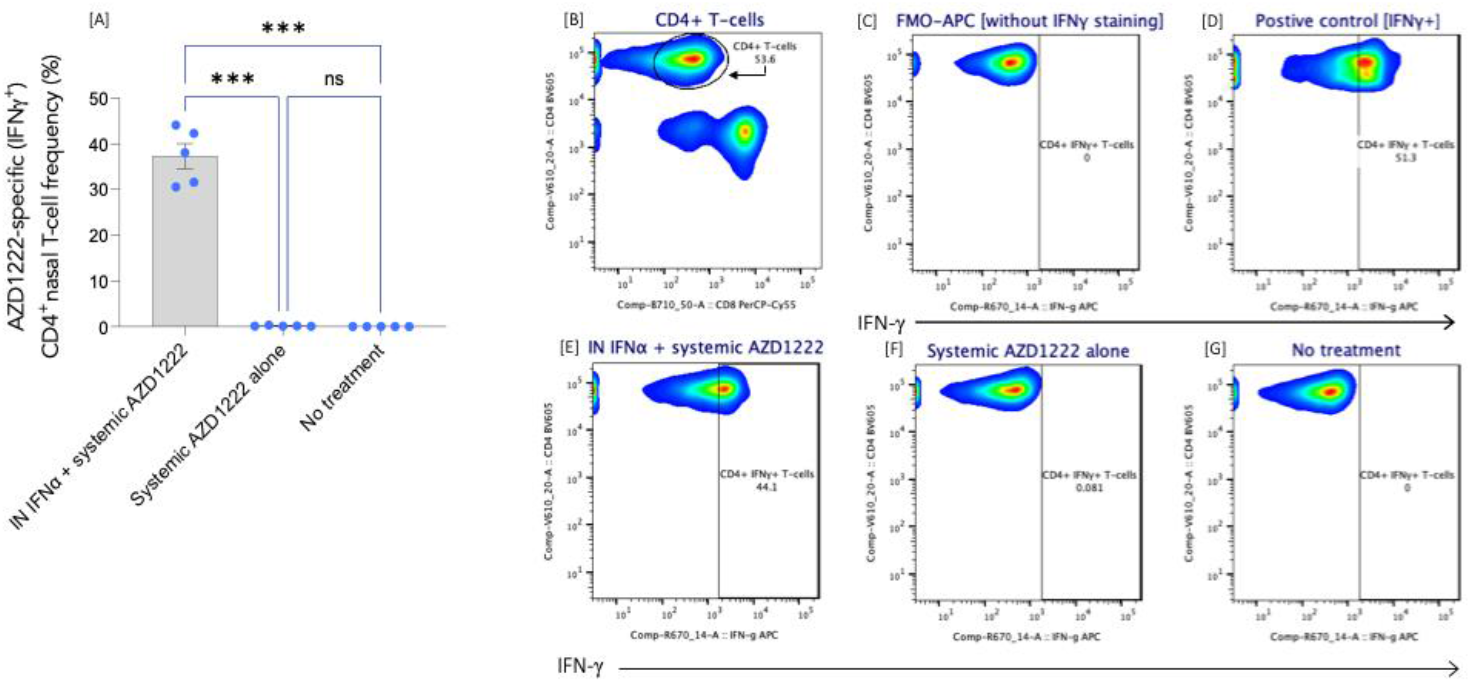
Intranasal IFN-α and systemic AZD1222 coadministration induced vaccine-specific CD4^+^ T-cell responses in the URT. [A] IFN-γ expression in CD4^+^ T-cells was significantly higher in the intranasal IFNα + systemic AZD1222 group compared to systemic AZD1222 alone and also compared to the untreated group. Data were analysed by applying two-way ANOVA and post hoc Tukey’s multiple comparison tests: \*\*\**P* < 0.001; ns: not significant. Representative flow cytometry plots show [B] gated CD4^+^ T-cells; and IFN-γ^+^CD4^+^ T-cell frequency in [C] FMO negative control, without IFN-γ–APC staining; [D] positive control; [E] intranasal IFN-α + systemic AZD1222 group; [F] systemic AZD1222 alone group; [G] no treatment group. IFN-γ expression is represented along the y-axis.

**Figure 2.**
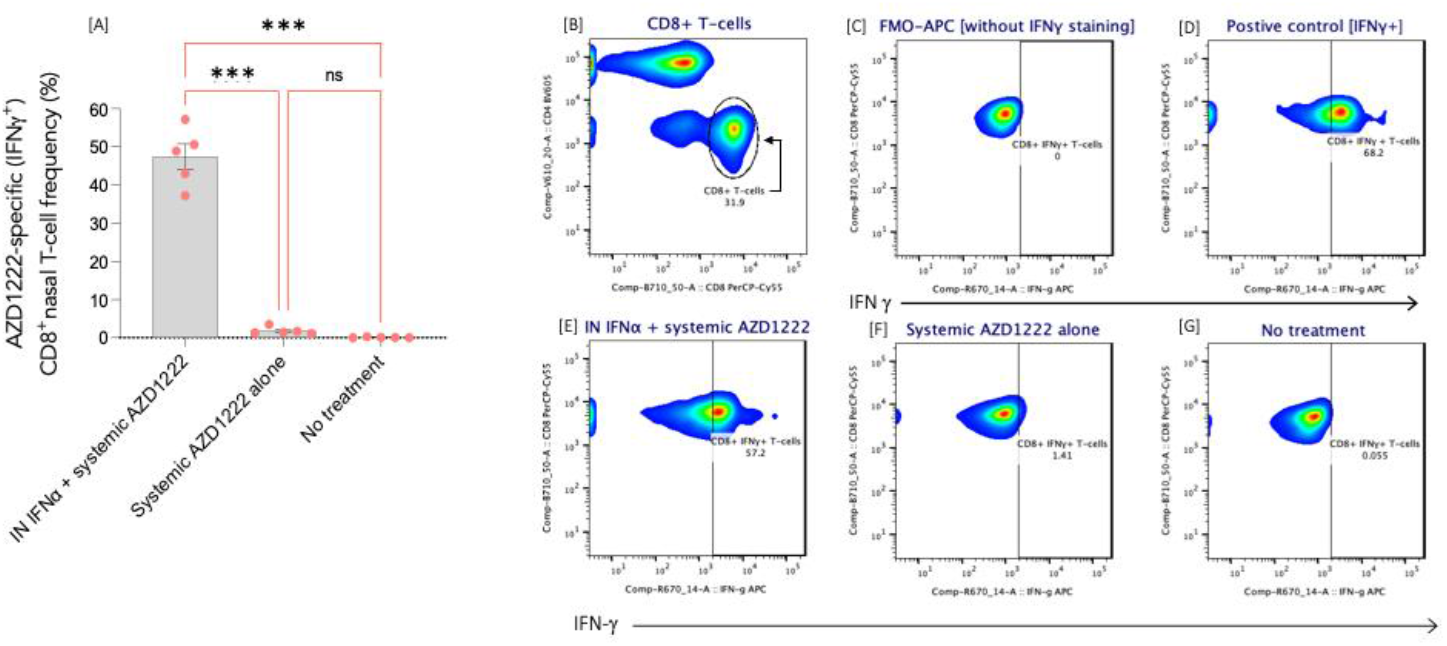
Intranasal IFN-α and systemic AZD1222 coadministration induced vaccine-specific CD8^+^ T-cell responses in the URT. [A] IFN-γ expression in CD8^+^ T-cells was significantly higher in the intranasal IFNα + systemic AZD1222 group compared to systemic AZD1222 alone and also compared to the untreated group. Data were analysed by applying two-way ANOVA and post hoc Tukey’s multiple comparison tests: \*\*\**P* < 0.001; ns: not significant. Representative flow cytometry plots show [B] CD8^+^ T-cells; and IFN-γ^+^CD8^+^ T-cell frequency in [C] FMO negative control, without IFN-γ–APC staining; [D] positive control; [E] intranasal IFN-α + systemic AZD1222 group; [F] systemic AZD1222 alone group; [G] no treatment group. IFN-γ expression is represented along the y-axis.

### Intranasal IFN-α administration alone does not induce T-cell activation

To determine whether IFN-α alone could induce T-cell activation in the URT, mice were administered intranasal IFN-α alone, without coadministration of (systemic) AZD1222. Using identical gating and *ex vivo* peptide stimulation strategies as described in Figure 1, no IFN-γ expression was detected in either CD4^+^ or CD8^+^ T-cells following stimulation with SARS-CoV-2 Spike peptides. As before, the percentage of IFN-γ^+^CD4^+^ or IFN-γ^+^CD8^+^ T-cells in peptide-stimulated samples was calculated after subtracting background signal from the negative control (FMO), and values were expressed as relative to the response observed in the positive control. FlowJo-based quantification determined that T-cell activation was undetectable at both 48 hours and 21 days post-IFN-α administration (Figure 3). The 48-hour and 21-day timepoints were chosen to capture potential immediate and/or transient effects (within 48 hours), and to align with the vaccination schedule (the 21-day time-point corresponding to the timing of the second vaccine dose with or without intranasal IFN-α coadministration). These observations confirm that administration of intranasal IFN-α alone is insufficient to trigger T-cell activation responses, in the absence of antigenic input from systemic vaccination.

**Figure 3.**
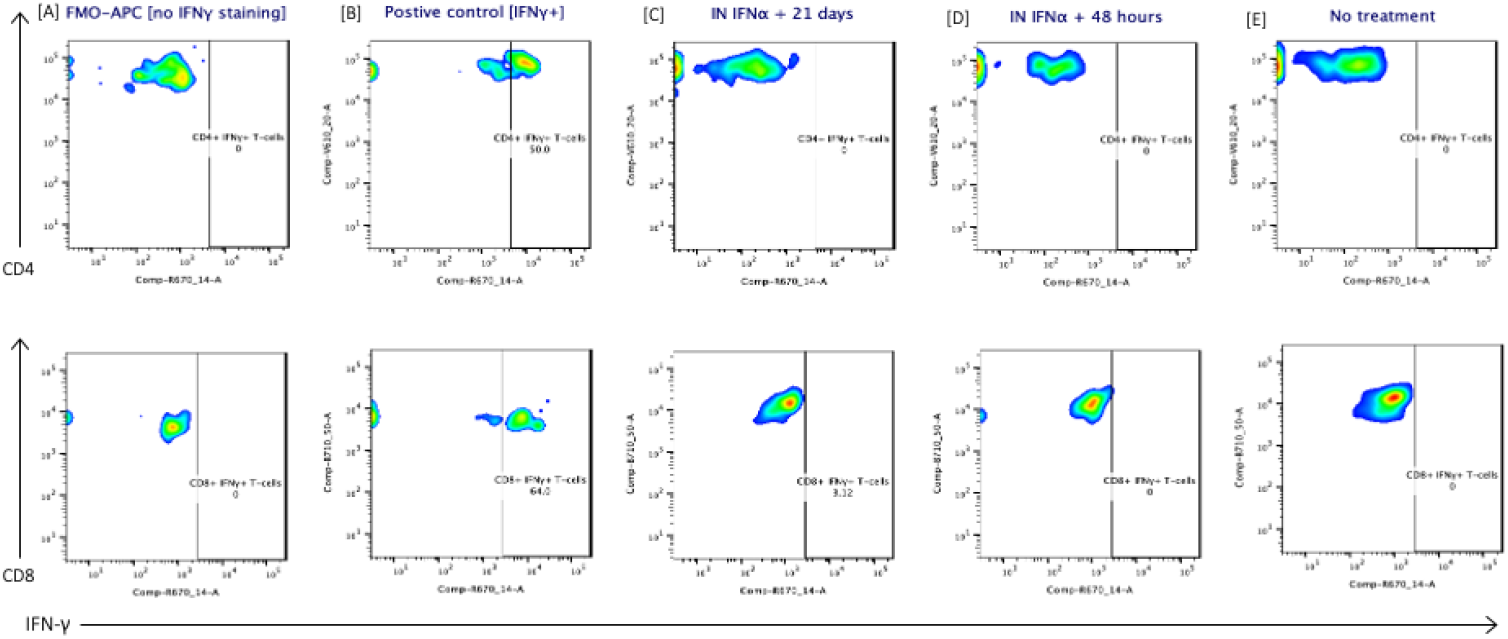
No CD4+ or CD8+ T-cell activation detected following intranasal IFN-α administration alone. Representative flow cytometric analysis for IFN-γ^+^CD4^+^ T-cells at 48 hours and 21 days post-IFN-α administration. [A] FMO negative control, without IFN-γ–APC staining; [B] positive control; [C] intranasal IFN-α + 21 days; [D] intranasal IFN-α + 48 hours; [E] no treatment. Representative flow cytometric analysis for IFN-γ^+^CD4^+^ T-cells are shown in the top row and IFN-γ^+^CD8^+^ T-cells are shown in the bottom row. CD4 or CD8 expression are represented in the x-axis and IFN-γ expression is represented along the y-axis.

## Discussion

In the current study, we demonstrate a novel vaccination strategy in which coadministration of intranasal IFN-α with simultaneous systemic SARS-CoV-2 vaccination induces vaccine-specific T-cell responses in the URT, despite the absence of local vaccine antigen. While intranasal vaccines have been investigated for decades, our approach circumvents the longstanding technical challenges associated with intranasal antigen delivery, by using innate immune modulation to direct systemic immunity to the URT mucosa. To our knowledge, this is the first report of antigen-independent recruitment of vaccine-specific T-cells to the URT upon concomitant systemic vaccination.

This strategy mimics the early host innate response to SARS-CoV-2 infection ^8^, and provides a novel means of establishing mucosal immunity at the primary site of viral entry and transmission. The URT, where host cells are initially targeted by respiratory pathogens, represents the main port of viral entry and exit and thus plays a central role in initial viral replication and onward transmission ^9,10^. When vaccines prevent transmission, this can in turn limit viral evolution towards highly transmissible immune-evasive variants ^11^. It is therefore anticipated that by directing vaccine-induced immunity to the site of pathogen entry at the URT, our innovation potentially offers a powerful strategy to interrupt transmission, slow down viral evolution and variant emergence, thereby enhancing population-level protection and protecting vulnerable populations.

Type 1 interferons (IFN-1s; which include IFN-α), are rapidly produced at mucosal surfaces in response to viral antigenic presence and drive expression of interferon-stimulated genes that inhibit viral replication and establish proinflammatory tissue conditions ^12-14^. Intranasal administration of exogenous IFN-α can replicate these effects even in the absence of infection, by engaging receptors IFNAR on epithelial and myeloid cells, and subsequently triggering a self-amplifying cascade of IFN-I production ^15,16^. The resulting local inflammatory milieu promotes the expression of chemokines, cytokines and adhesion molecules to amplify inflammation, and upregulate endothelial adhesion molecule expression in local vascular beds to facilitate transendothelial lymphocyte migration into the tissue ^12-14,17^. This chemotactic environment may thus enable the recruitment of vaccine-primed lymphocytes to the URT, even without local antigenic cues.

Importantly, there may also be value in capitalising on newly activated lymphocytes (such as those induced by systemic vaccination), which exhibit a transiently heightened capacity to migrate to inflamed tissues, due to upregulation of homing receptors (integrins, selectins, chemokine receptors) that increase sensitivity to chemotactic cues and enable trafficking to distant sites expressing inflammatory signals ^18,19^. For example, newly activated γδT17 cells have been shown to home to distant inflamed tissues, such as skin, to amplify local inflammation; while previously generated effector lymphocytes downregulate their homing receptors since they revert to a resting state, and exhibit less dynamic migration towards newly inflamed sites compared to freshly activated lymphocytes ^20-23^ thus reducing their responsiveness to inflammatory signals and limiting their potential for tissue residency. As such, IFN-α-mediated inflammation in the URT may preferentially recruit recently activated, vaccine-specific T-cells and promote their retention through the establishment of tissue-resident memory cells ^18^. It is hence plausible that newly activated lymphocytes in our model (induced by systemic AZD1222 vaccination) were recruited, from the lower respiratory tract or the circulation, to the URT, in response to local IFN-α signalling. Indeed, it has been demonstrated that while AZD1222 can induce lower respiratory tract mucosal responses, it does not elicit URT immunity.

In humans, intramuscular vaccination such as with AZD1222 (typically administered in the upper arm deltoid muscle) drains into the axillary lymph nodes, facilitating systemic priming and mobilisation of immune cells capable of homing to the lower respiratory tract ^1,4^. In contrast, murine intramuscular injections are typically administered in the hind limb, which drain to the inguinal and iliac nodes to generate systemic responses that do not typically extend to the respiratory tract ^24-26^. However, intraperitoneal administration may represent an appropriate proxy for human systemic vaccination in the context of modelling systemic and lower respiratory immunity. This route provides broad systemic priming via drainage to mediastinal lymph nodes and spleen ^27,28^, and can facilitate homing to the lower respiratory mucosa ^29^. Therefore, the intraperitoneal vaccination route employed in the current murine study may better approximate the immunological dynamics of human intramuscular vaccination than the standard murine intramuscular route.

SARS-CoV-2 continues to be responsible for significant morbidity and mortality, and vulnerable populations (particularly the immunocompromised and those at the extremes of age) remain disproportionately affected ^30-32^. These groups also face elevated risks of respiratory, cardiopulmonary, vascular and cognitive sequelae ^33^. Cyclical infection surges persist, driven by ongoing community transmission, viral evolution, and inadequate vaccine-induced protection against infection ^34-36^. High transmission rates in vaccinated populations can accelerate the emergence of vaccine-resistant variants with increased transmissibility and immune evasion ^11,36^. Persistent infections, particularly in immunocompromised individuals, may lead to tissue reservoirs where key mutations accumulate, contributing to both post-acute sequelae and variant emergence ^37,38^. Unless viral transmission is controlled, recurrent waves of infection and the emergence of new vaccine-resistant variants will likely continue for the foreseeable future. Our findings therefore offer a potential strategy to promote URT mucosal defences and aid in limiting the cycle of SARS-CoV-2 transmission and evolution. The strategy presented here with targeted innate stimulation at the URT mucosa with exogenous IFN-α, when coupled with systemic vaccination, may further be widely applicable to vaccines against diverse respiratory pathogens that are known to trigger or be modulated by IFN-α-mediated antiviral responses ^39-43^. This strategy also aligns with the WHO’s ‘Preparedness and Resilience for Emerging Threats’ initiative, which emphasises the need for adaptable, rapidly deployable pandemic responses ^44^. Thus, the proposed strategy represents a potentially powerful adjunct in public health emergencies and pandemic response toolkits, as well as in other scenarios where rapid roll-out is necessary, such as in low-resource settings. Further, high patient compliance is likely, with simultaneous administration of intranasal IFN-α with parenteral vaccines at clinics, providing a streamlined solution that minimises wastage and improves compliance. Notably, this approach could also reduce the need for frequent vaccine reformulation in response to evolving variants.

## Limitations of the study

While viral vector vaccines against SARS-CoV-2 such as AZD1222 are no longer widely used (mRNA vaccines dominate the seasonal SARS-CoV-2 vaccine updates in the global 7MM, with −80°C storage requirements), the adenoviral vector technology allows for vaccination strategies to be used that are less dependent on a cold chain. As such, testing our intervention using different vaccination platforms may be relevant for future pandemics or other applications requiring rapid deployment. Further studies are required to comprehensively characterise the local immune landscape in the URT following coadministration of intranasal IFN-α and a systemically delivered respiratory viral vaccine. In particular, nasal mucosal immunoprofiling and viral challenge studies may be valuable for defining immune composition and confirming protective efficacy, thereby informing subsequent translational evaluation prior to larger-scale trials.

## Author contributions

R.F. conceived and designed the study, performed experiments with assistance from the University of Edinburgh’s Bioresearch & Veterinary Services (BVS) and the IRR Flow Cytometry Facility, analysed the data and prepared the manuscript. D.H.D. and J.S. contributed intellectual input and critically appraised the work. D.H.D. was responsible for providing the space and equipment for this work to be conducted. All authors approved the final version of the manuscript.

## Acknowledgements

The authors would like acknowledge the support of both AstraZeneca and Sarah Gilbert, University of Oxford, for providing AZD1222 vaccine doses for this work to be conducted, and we are grateful to AstraZeneca for approving the final version of the manuscript. We are also very grateful to Edinburgh Innovations (Jenny Cameron, Andrew McBride and Mark Craighead) for establishing connections with commercial partners; William Mungal and Rosie Shield at BVS for performing vaccinations, administering intranasal IFN-α treatments, monitoring animals, and harvesting tissue; the IRR Flow Cytometry Facility for support with sample acquisition; and members of the D.H.D. and J.S. research groups for their assistance with laboratory logistics.

## References

1. van Doremalen, N., Lambe, T., Spencer, A., Belij-Rammerstorfer, S., Purushotham, J.N., Port, J.R., Avanzato, V.A., Bushmaker, T., Flaxman, A., Ulaszewska, M., et al. (2020). ChAdOx1 nCoV-19 vaccine prevents SARS-CoV-2 pneumonia in rhesus macaques. Nature 586, 578–582. 10.1038/s41586-020-2608-y.

2. Tang, J., Zeng, C., Cox, T.M., Li, C., Son, Y.M., Cheon, I.S., Wu, Y., Behl, S., Taylor, J.J., Chakraborty, R., et al. (2022). Respiratory mucosal immunity against SARS-CoV-2 following mRNA vaccination. Sci Immunol, eadd4853. 10.1126/sciimmunol.add4853.

3. Lim, J.M.E., Tan, A.T., Le Bert, N., Hang, S.K., Low, J.G.H., and Bertoletti, A. (2022). SARS-CoV-2 breakthrough infection in vaccinees induces virus-specific nasal-resident CD8+ and CD4+ T cells of broad specificity. J Exp Med 219. 10.1084/jem.20220780.

4. Fraser, R., Orta-Resendiz, A., Mazein, A., & Dockrell, D. H. (2023). Upper respiratory tract mucosal immunity for SARS-CoV-2 vaccines. Trends in Molecular Medicine, Advance online publication. 10.1016/j.molmed.2023.01.003.

5. Brokstad, K.A., Eriksson, J.C., Cox, R.J., Tynning, T., Olofsson, J., Jonsson, R., and Davidsson, A. (2002). Parenteral vaccination against influenza does not induce a local antigen-specific immune response in the nasal mucosa. J Infect Dis 185, 878–884. 10.1086/339710.

6. Mao, T., Israelow, B., Peña-Hernández, M.A., Suberi, A., Zhou, L., Luyten, S., Reschke, M., Dong, H., Homer, R.J., Saltzman, W.M., and Iwasaki, A. (2022). Unadjuvanted intranasal spike vaccine elicits protective mucosal immunity against sarbecoviruses. Science 378, eabo2523. 10.1126/science.abo2523.

7. Tregoning, J.S., Buffa, V., Oszmiana, A., Klein, K., Walters, A.A., and Shattock, R.J. (2013). A “prime-pull” vaccine strategy has a modest effect on local and systemic antibody responses to HIV gp140 in mice. PloS one 8, e80559. 10.1371/journal.pone.0080559.

8. Cheemarla, N.R., Watkins, T.A., Mihaylova, V.T., Wang, B., Zhao, D., Wang, G., Landry, M.L., and Foxman, E.F. (2021). Dynamic innate immune response determines susceptibility to SARS-CoV-2 infection and early replication kinetics. J Exp Med 218. 10.1084/jem.20210583.

9. Loske, J., Röhmel, J., Lukassen, S., Stricker, S., Magalhães, V.G., Liebig, J., Chua, R.L., Thürmann, L., Messingschlager, M., Seegebarth, A., et al. (2022). Pre-activated antiviral innate immunity in the upper airways controls early SARS-CoV-2 infection in children. Nature Biotechnology 40, 319–324. 10.1038/s41587-021-01037-9.

10. Nakayama, T., Lee, I.T., Jiang, S., Matter, M.S., Yan, C.H., Overdevest, J.B., Wu, C.-T., Goltsev, Y., Shih, L.-C., Liao, C.-K., et al. (2021). Determinants of SARS-CoV-2 entry and replication in airway mucosal tissue and susceptibility in smokers. Cell Reports Medicine 2, 100421. 10.1016/j.xcrm.2021.100421.

11. Read, A.F., Baigent, S.J., Powers, C., Kgosana, L.B., Blackwell, L., Smith, L.P., Kennedy, D.A., Walkden-Brown, S.W., and Nair, V.K. (2015). Imperfect Vaccination Can Enhance the Transmission of Highly Virulent Pathogens. PLoS Biol 13, e1002198. 10.1371/journal.pbio.1002198.

12. Muller, W.A. (2009). Mechanisms of transendothelial migration of leukocytes. Circ Res 105, 223–230. 10.1161/circresaha.109.200717.

13. Groom, J.R., and Luster, A.D. (2011). CXCR3 ligands: redundant, collaborative and antagonistic functions. Immunol Cell Biol 89, 207–215. 10.1038/icb.2010.158.

14. Luft, T., Pang, K.C., Thomas, E., Hertzog, P., Hart, D.N., Trapani, J., and Cebon, J. (1998). Type I IFNs enhance the terminal differentiation of dendritic cells. J Immunol 161, 1947–1953.

15. Raftery, N., and Stevenson, N.J. (2017). Advances in anti-viral immune defence: revealing the importance of the IFN JAK/STAT pathway. Cell Mol Life Sci 74, 2525–2535. 10.1007/s00018-017-2520-2.

16. Rahmatpanah, F., Agrawal, S., Jaiswal, N., Nguyen, H.M., McClelland, M., and Agrawal, A. (2019). Airway epithelial cells prime plasmacytoid dendritic cells to respond to pathogens via secretion of growth factors. Mucosal Immunol 12, 77–84. 10.1038/s41385-018-0097-1.

17. Montoya, M., Schiavoni, G., Mattei, F., Gresser, I., Belardelli, F., Borrow, P., and Tough, D.F. (2002). Type I interferons produced by dendritic cells promote their phenotypic and functional activation. Blood 99, 3263–3271. 10.1182/blood.v99.9.3263.

18. Mackay, L.K., Rahimpour, A., Ma, J.Z., Collins, N., Stock, A.T., Hafon, M.L., Vega-Ramos, J., Lauzurica, P., Mueller, S.N., Stefanovic, T., et al. (2013). The developmental pathway for CD103(+)CD8+ tissue-resident memory T cells of skin. Nat Immunol 14, 1294–1301. 10.1038/ni.2744.

19. Hickman, H.D., Li, L., Reynoso, G.V., Rubin, E.J., Skon, C.N., Mays, J.W., Gibbs, J., Schwartz, O., Bennink, J.R., and Yewdell, J.W. (2011). Chemokines control naive CD8+ T cell selection of optimal lymph node antigen presenting cells. J Exp Med 208, 2511–2524. 10.1084/jem.20102545.

20. McKenzie, D.R., Kara, E.E., Bastow, C.R., Tyllis, T.S., Fenix, K.A., Gregor, C.E., Wilson, J.J., Babb, R., Paton, J.C., Kallies, A., et al. (2017). IL-17-producing γδ T cells switch migratory patterns between resting and activated states. Nat Commun 8, 15632. 10.1038/ncomms15632.

21. Ramírez-Valle, F., Gray, E.E., and Cyster, J.G. (2015). Inflammation induces dermal Vγ4+ γδT17 memory-like cells that travel to distant skin and accelerate secondary IL-17-driven responses. Proc Natl Acad Sci U S A 112, 8046–8051. 10.1073/pnas.1508990112.

22. Sallusto, F., Lenig, D., Förster, R., Lipp, M., and Lanzavecchia, A. (1999). Two subsets of memory T lymphocytes with distinct homing potentials and effector functions. Nature 401, 708–712. 10.1038/44385.

23. Skon, C.N., Lee, J.Y., Anderson, K.G., Masopust, D., Hogquist, K.A., and Jameson, S.C. (2013). Transcriptional downregulation of S1pr1 is required for the establishment of resident memory CD8+ T cells. Nat Immunol 14, 1285–1293. 10.1038/ni.2745.

24. Qiu, J., Peng, S., Yang, A., Ma, Y., Han, L., Cheng, M.A., Farmer, E., Hung, C.F., and Wu, T.C. (2018). Intramuscular vaccination targeting mucosal tumor draining lymph node enhances integrins-mediated CD8+ T cell infiltration to control mucosal tumor growth. Oncoimmunology 7, e1463946. 10.1080/2162402x.2018.1463946.

25. Schutsky, K., Curtis, D., Bongiorno, E.K., Barkhouse, D.A., Kean, R.B., Dietzschold, B., Hooper, D.C., and Faber, M. (2013). Intramuscular inoculation of mice with the live-attenuated recombinant rabies virus TriGAS results in a transient infection of the draining lymph nodes and a robust, long-lasting protective immune response against rabies. J Virol 87, 1834–1841. 10.1128/jvi.02589-12.

26. Nakajima, Y., Asano, K., Mukai, K., Urai, T., Okuwa, M., Sugama, J., and Nakatani, T. (2018). Near-Infrared Fluorescence Imaging Directly Visualizes Lymphatic Drainage Pathways and Connections between Superficial and Deep Lymphatic Systems in the Mouse Hindlimb. Sci Rep 8, 7078. 10.1038/s41598-018-25383-y.

27. Sheasley-O’Neill, S.L., Brinkman, C.C., Ferguson, A.R., Dispenza, M.C., and Engelhard, V.H. (2007). Dendritic cell immunization route determines integrin expression and lymphoid and nonlymphoid tissue distribution of CD8 T cells. J Immunol 178, 1512–1522. 10.4049/jimmunol.178.3.1512.

28. Liu, M., Silva-Sanchez, A., Randall, T.D., and Meza-Perez, S. (2021). Specialized immune responses in the peritoneal cavity and omentum. J Leukoc Biol 109, 717–729. 10.1002/jlb.5mir0720-271rr.

29. Jearanaiwitayakul, T., Apichirapokey, S., Chawengkirttikul, R., Limthongkul, J., Seesen, M., Jakaew, P., Trisiriwanich, S., Sapsutthipas, S., Sunintaboon, P., and Ubol, S. (2021). Peritoneal Administration of a Subunit Vaccine Encapsulated in a Nanodelivery System Not Only Augments Systemic Responses against SARS-CoV-2 but Also Stimulates Responses in the Respiratory Tract. Viruses 13. 10.3390/v13112202.

30. Chakraborty, C., Bhattacharya, M., and Lee, S.S. (2025). Excess mortality in older adults and cumulative excess mortality across all ages during the COVID-19 pandemic in the 20 countries with the highest mortality rates worldwide. Osong Public Health Res Perspect 16, 42–58. 10.24171/j.phrp.2024.0186.

31. Lippi, G., and Sanchis-Gomar, F. (2025). Mortality of Post-COVID-19 Condition: 2025 Update. COVID 5, 11.

32. Spinner, C.D., Bell, S., Einsele, H., Tremblay, C., Goldman, M., Chagla, Z., Finckh, A., Edwards, C.J., Aurer, I., Launay, O., et al. (2025). Is COVID-19 Still a Threat? An Expert Opinion Review on the Continued Healthcare Burden in Immunocompromised Individuals. Adv Ther 42, 666–719. 10.1007/s12325-024-03043-0.

33. CDC (2025). Underlying Conditions and the Higher Risk for Severe COVID-19: https://www.cdc.gov/covid/hcp/clinical-care/underlying-conditions.html.

34. Ghafari, M., Hall, M., Golubchik, T., Ayoubkhani, D., House, T., MacIntyre-Cockett, G., Fryer, H.R., Thomson, L., Nurtay, A., Kemp, S.A., et al. (2024). Prevalence of persistent SARS-CoV-2 in a large community surveillance study. Nature 626, 1094–1101. 10.1038/s41586-024-07029-4.

35. Venkatakrishnan, A.J., Anand, P., Lenehan, P., Ghosh, P., Suratekar, R., Siroha, A., Chowdhury, D.R., O’Horo, J.C., Yao, J.D., Pritt, B.S., et al. (2021). Antigenic minimalism of SARS-CoV-2 is linked to surges in COVID-19 community transmission and vaccine breakthrough infections. medRxiv, 2021.2005.2023.21257668. 10.1101/2021.05.23.21257668.

36. Rella, S.A., Kulikova, Y.A., Dermitzakis, E.T., and Kondrashov, F.A. (2021). Rates of SARS-CoV-2 transmission and vaccination impact the fate of vaccine-resistant strains. Sci Rep 11, 15729. 10.1038/s41598-021-95025-3.

37. Swank, Z., Senussi, Y., Manickas-Hill, Z., Yu, X.G., Li, J.Z., Alter, G., and Walt, D.R. (2023). Persistent Circulating Severe Acute Respiratory Syndrome Coronavirus 2 Spike Is Associated With Post-acute Coronavirus Disease 2019 Sequelae. Clin Infect Dis 76, e487–e490. 10.1093/cid/ciac722.

38. Gonzalez-Reiche, A.S., Alshammary, H., Schaefer, S., Patel, G., Polanco, J., Carreño, J.M., Amoako, A.A., Rooker, A., Cognigni, C., Floda, D., et al. (2023). Sequential intrahost evolution and onward transmission of SARS-CoV-2 variants. Nat Commun 14, 3235. 10.1038/s41467-023-38867-x.

39. Guerrero-Plata, A., Baron, S., Poast, J.S., Adegboyega, P.A., Casola, A., and Garofalo, R.P. (2005). Activity and regulation of alpha interferon in respiratory syncytial virus and human metapneumovirus experimental infections. J Virol 79, 10190–10199. 10.1128/jvi.79.16.10190-10199.2005.

40. Matzinger, S.R., Carroll, T.D., Dutra, J.C., Ma, Z.M., and Miller, C.J. (2013). Myxovirus resistance gene A (MxA) expression suppresses influenza A virus replication in alpha interferon-treated primate cells. J Virol 87, 1150–1158. 10.1128/jvi.02271-12.

41. Dieterich, C., and Relman, D.A. (2011). Modulation of the host interferon response and ISGylation pathway by B. pertussis filamentous hemagglutinin. PLoS One 6, e27535. 10.1371/journal.pone.0027535.

42. Boeren, M., Van Breedam, E., Buyle-Huybrecht, T., Lebrun, M., Meysman, P., Sadzot-Delvaux, C., Van Tendeloo, V.F., Mortier, G., Laukens, K., Ogunjimi, B., et al. (2022). Activation of Interferon-Stimulated Genes following Varicella-Zoster Virus Infection in a Human iPSC-Derived Neuronal In Vitro Model Depends on Exogenous Interferon-α. Viruses 14. 10.3390/v14112517.

43. Hernandez, N., Bucciol, G., Moens, L., Le Pen, J., Shahrooei, M., Goudouris, E., Shirkani, A., Changi-Ashtiani, M., Rokni-Zadeh, H., Sayar, E.H., et al. (2019). Inherited IFNAR1 deficiency in otherwise healthy patients with adverse reaction to measles and yellow fever live vaccines. J Exp Med 216, 2057–2070. 10.1084/jem.20182295.

44. WHO (2023). Preparedness and Resilience for Emerging Threats (PRET).

